# Nucleus of the lateral olfactory tract (NLOT): a hub linking water homeostasis-associated SON-AVP circuit and neocortical regions to promote social behavior under osmotic challenge

**DOI:** 10.1101/2022.07.01.498472

**Authors:** Oscar R. Hernández-Pérez, Vito S. Hernández, Mario A. Zetter, Lee E. Eiden, Limei Zhang

## Abstract

Homeostatic challenges increase the drive for social interaction. The neural activity that prompts this motivation remains poorly understood. Here, we identify direct projections from the hypothalamic supraoptic nucleus (SON) to the cortico-amygdalar nucleus of the lateral olfactory tract (NLOT). Dual *in situ* hybridization (DISH) with probes for PACAP, and VGLUT1, VGLUT2, V1a and V1b revealed a population of vasopressin-receptive PACAPergic neurons in NLOT layer 2 (NLOT2). Water deprivation (48 hours, WD48) increased sociability compared to euhydrated subjects, assessed with the three-chamber social interaction test (3CST). Fos expression immunohistochemistry showed NLOT and its main efferent regions had further increases in rats subjected to WD48+3CST. These regions strongly expressed PAC1 mRNA. Microinjections of AVP into NLOT produced similar changes in sociability to water deprivation, and these were reduced by co-injection of V1a or V1b antagonists along with AVP. We conclude that during challenge to water homeostasis, there is a recruitment of a glutamatergic-multi-peptidergic cooperative circuit that promotes social behavior.

## 1. INTRODUCTION

Neuropeptides acting as neurotransmitters in the brain participate in circuits which monitor multiple environmental inputs, and access brain state-related information, in order to produce highly integrated responses that are appropriately tuned to the specifics of a given situation requiring motor decision-making (action)^(1)^. An area of research relevant to the general question of how multiple-circuit integration leads to specific behaviors occurs, is the study of the microanatomy, neurochemistry, and cellular and behavioral functions of vasopressinergic magnocellular neurosecretory neurons (AVPMNNs) of the mammalian brain. This is because AVPMNNs have very well-defined roles as neurosecretory neurons that release vasopressin into the bloodstream for hormonally-driven osmotic regulation (hydromineral *homeostasis*), and as well, via dual projections within the brain parenchyma, release vasopressin as a neurotransmitter to affect limbic, hypothalamic, and other brain circuits that participate in behavioral prioritization responsive to competing homeostatic drives, i.e. *allostasis* ^(2)^.

In previous studies, we have demonstrated that activation of the AVPMNN system in the hypothalamic paraventricular nucleus (PVN) by alteration of osmotic balance through water deprivation or salt loading activates both limbs of this dually-projecting system. This results not only in changes in blood levels of vasopressin indicative of hormonal homeostatic regulation, but also in changes in electrical activity and even synaptic protein expression at vasopressinergic terminal field regions of the hippocampus, amygdala, locus coeruleus and other locations. These are indicative of potentially profound effects on behaviors ^(3-9)^ and thus linking the original homeostatic osmoregulatory drive, to allostatic drives that must act in concert with it.

Social behavior is one of the core factors facilitating an individual’s chances of survival. However, the neural substrates underlying this behavioral characteristic remain poorly understood. Vasopressin has been shown to participate in several aspects of social behavior; for instance, mice deficient for V1a and V1b receptors show social deficits^(10-12)^. Brattleboro rats lacking AVP expression have reduced social interaction ^(13, 14)^, and altered AVP transmission has been described in pathologies characterized by social deficits ^(15, 16)^.

In contrast to the PVN, the hypothalamic supraoptic nucleus (SON), which contains another major population of AVPMNNs, responsible for controlling body water balance, has been less investigated regarding ascending projections, as well as concerning its roles in behavioral adaptation. This lack of information is mainly due to its location at the base of the brain, adjacent to the optic chiasm. Using *in vivo* electrophysiological recording, Inyushkin et al. ^(17)^ demonstrated a clear dual projection system, neurohypophysial and central, from SON AVPMNNs, and suggested the existence of projections from these neurons in the SON that are much more widespread and longer than had previously been suspected. SON’s innervation to dorsal and ventral hippocampus and central amygdala was subsequently demonstrated using the fluorogold method ^(6, 18)^.

The nucleus of the lateral olfactory tract (NLOT) is located in the anterior cortico-amygdalar region adjacent to the ventral surface of the brain. This structure is connected with the main olfactory bulb, and the piriform and insular cortices and is implicated in non-pheromonal behavior such as feeding ^(19)^. It has been described as a three-layered structure^(20)^. Its heterogeneous neuronal composition suggests different neuroepithelial origins of the cells that populate its three layers ^(21)^.

The aim of the present study was to evaluate whether the NLOT containing vasopressin-responsive neurons is important in mediating the social effects of AVP. By analyzing Golgi-Cox staining, we observed direct projections from SON to the nucleus of lateral olfactory tract (NLOT) of the cortico-amygdalar complex. Fluoro-Gold injection into the NLOT revealed retrogradely labeled AVP-positive somata in SON. We previously reported that the main neuronal population of the NLOT co-expresses the neuropeptide PACAP as well as vesicular glutamate transporters 1 and 2 (VGLUT1 and VGLUT2). Here, using the dual in situ hybridization method, we demonstrate that the principal cell population PACAP/VGLUT1/VGLUT2 co-express vasopressin receptors V1a and V1b. Considering these observations, we devised a behavioral experiment using 48 hours of water deprivation (WD48) and the three-chamber social interaction test (3CST) ^(22, 23)^, designed to quantify the levels of sociability, defined as the tendency to approach and remain proximal to an unfamiliar conspecific (we called “rat *stranger*” hereafter). WD48 significantly increased the sociability parameters. Fos expression assessment showed that the downstream regions of AVPMNNs and PACAP-NLOT had further increases in rats subjected to WD48+3CSI. The downstream regions of PACAP-NLOT strongly expressed PAC1 mRNA. AVP and NLOT involvement in this increased sociability and Fos expression was further demonstrated through microinjections of AVP, AVP+V1a or V1b antagonists targeting NLOT. The AVP microinjection alone produced a similar increase in sociability and Fos expression as that produced by WD48, but if applied together with V1a or V1b antagonists, the AVP-stimulated increases were ablated. These results suggest that under situations where homeostasis is compromised by osmotic challenge, there is a recruitment of a glutamatergic-multi-peptidergic cooperative circuit that is able to promote behavioral adaptation through social interaction.

## 2. EXPERIMENTAL PROCEDURES

### 2.1 Animals

One hundred and forty-four adult male Wistar rats of 280 ± 20 g were obtained from the local animal facility. Rats were housed four per cage in a controlled environment (temperature 24 ºC and illumination 12 h/12 h (lights on at 7:00 to 19:00 h) with water and food *ad libitum*). All animal procedures were approved by the *Comision de Investigacion y Etica de la Facultad de Medicina, Universidad Nacional Autonoma de Mexico* (approval number: CIEFM-062-2016). Efforts were taken to minimize animal suffering throughout all experimental procedures.

### 2.2 Social behavior assessment under water deprivation

A first experiment was devised to assess the effects of water deprivation on social behavior. Twenty-six rats were randomly assigned to 48h of water deprivation (WD48, n=14) or water *ad libitum* (control, n=12). The rationale for this manipulation was that WD48 up-regulates the metabolic activity of the hypothalamic AVPMNNs with only a modest increase in plasma osmolarity ^(5, 24)^. Social behavior was tested in the three-chamber social interaction test (3CST). The test was carried out essentially as described elsewhere ^(22, 23)^. The test was conducted in a three-chambered box made of acrylic (100 cm x 30 cm x 30 cm), the central chamber (20 cm x 30 cm) and two distal chambers (40 cm x 30 cm) each. Chambers were connected via open doors (10 cm x 10 cm). One day before the experiment, the rats were allowed to explore the apparatus freely for 10 min to be habituated to the experimental device. On the day of the experiment, an empty cage was placed in a compartment (*non-social* compartment), a similar cage containing a male rat *stranger* was placed in the opposite chamber (*social* compartment). The experiment started by putting the experimental subjects in the central chamber of the apparatus and allowing access to all chambers for 10 minutes. The time the rat spent in the *social* chamber and the number of approaches/sniffing to the rat *stranger* were scored.

### 2.3 Immunohistochemistry and immunofluorescence

Rats were deeply anesthetized with sodium pentobarbital (63 mg/kg, Sedalpharma, México) and perfused transaortically with 0.9% NaCl followed by cold fixative containing 4% paraformaldehyde in 0.1 M sodium phosphate buffer (PB, pH 7.4) plus 15% v/v saturated picric acid solution for 15 min. Brains were immediately removed, blocked, thoroughly rinsed with PB 0.1M, and sectioned at 70 µm from forebrain to hindbrain using a Leica VT 1000S vibratome. Sections were blocked with 20% normal donkey serum in Tris-buffered (0.05 M, pH 7.4), NaCl (0.9%), plus 0.3% of Triton X-100 (TBST) for 1h at room temperature, immunoreacted overnight with rabbit anti-Fos primary antibody (0.1µg/ml, Santa Cruz Biotechnology, SC-52, CA, USA) or rabbit anti vasopressin primary antibody (1:2000 dilution, kind gift from Professor Ruud Buijs), washed and incubated with secondary biotinylated goat anti-rabbit antibody (3 µg/ml, Vector Labs, BA1000, CA, USA) followed by incubation in Vectastain Elite ABC kit solution (Vector Labs, Burlingame, CA, USA) and detection with a DAB-peroxidase reaction. For Immunofluorescent processing we used a donkey anti rabbit Alexa Fluor 594 secondary antibody (4 µg/ml, Thermo Fisher, A-21207, Massachusetts, USA). Some photomicrographs were presented digitally inverted (negative mode) to enhance the visibility of DAB-labelled AVP fibers.

### 2.4 Neuronal Activation assessment after 3CST

To assess the pattern of neuronal activation induced in the brain by the 3CST and/or water deprivation, we formed four groups of n=5 rats: Control (same conditions as other groups but undisturbed until perfusion time); Social interaction (rats that were under ad-libitum water access until the 3CST); 48h WD (rats deprived of water during the 48h previous to the perfusion with no other disturbance) and 48h WD + social Interaction (rats that underwent 48h of water deprivation before the 3CST experiment). Rats evaluated by the 3CST were perfused 60 min after finishing the behavioral test; the other groups were perfused at the same circadian time. After perfusion immunohistochemistry against Fos was performed as above mentioned.

Regions of interest were identified using a low magnification objective referencing with the rat brain stereotaxic atlas ^(25)^. The number of Fos+ nuclei in each identified region was quantified under a 40x objective (0.196 mm2). A table comparing the four groups was constructed with semi-quantitative criteria. i.e., “+”: 1 to 25 per field; “++”: 26 to 50 per field; “+++”: 51 to 75 per field; “++++”: 76 to 100 per field; “+++++”: >100 per field (field area: 0.196 mm2, for simplicity we used 0.2 mm^2^).

### 2.5 Fluorogold retrograde tracing

The Fluorogold retrograde method was performed as previously reported ^(6, 18)^. Rats (male, n=4, 280-300g) were deeply anaesthetized using a 1:1 mixture of xylazine (20 mg/ml, Procin, Mexico) and ketamine (100 mg/ml, Inoketam, Virbac, Mexico) administered a dose of 1 ml/kg body weight intraperitoneally. Rats were fixed in a stereotaxic frame, and the retrograde tracer Fluoro-Gold (FG, Fluorochrome, LLC, Denver, Colorado 80218, USA), dissolved to a concentration of 1% in 0.1 M cacodylate buffer (pH 7.5), was delivered into the nucleus of the lateral olfactory tract (NLOT) using a glass micropipette with an inner tip diameter of around 40 µm. Current used for iontophoresis was 0.1 µA with a 5-s pulse duration and a 50% duty cycle for 20 min. The coordinates for positioning the pipette in NLOT were: AP -1.4 mm, ML ± 3.2 mm from bregma, and DV -9.4 mm from the skull surface, according to a stereotaxic atlas ^(25)^. An additional time-lapse of 10 min was allowed to prevent backflow of tracer up the injection track. Before recovering from anesthesia, rats received 0.4 mg/kg i.p. ketorolac (Apotex, Mexico) and 50 mg/kg i.p. ceftriaxone (Kendric, Mexico) to reduce pain and risk of infection. This therapeutic scheme was repeated 1/24 hr. for three consecutive days. Three weeks after the FG injections, the rats were perfused (*vide supra*). Free floating coronal sections were obtained with a vibratome. AVP immunofluorescence reaction was made using rabbit anti-AVP antibody as described above. Observations were made using a Zeiss LSM 880 confocal microscope. FG was observed by using a UV excitation filter, and images were artificially assigned a green color for better visualization.

### 2.6 Golgi-Cox impregnation and neuronal 2-D reconstruction

For the argentic impregnation, four 300g male rats were deeply anesthetized and decapitated. Coronal blocks approximately 5 mm thick, containing the hypothalamus, were cut with a sharp blade and briefly rinsed with PB 0.1M. Blocks were immersed in sequential impregnation A/B and C solutions as indicated in the FD Rapid GolgiStain Kit (FD Neurotechnologies, Ellicott City, MD), during the following two weeks, after which 150 µm coronal sections containing the NLOT and SON were obtained using a vibratome with the cutting chamber filled with solution C, and mounted on gelatin-coated microscope slides, dried overnight at room temperature in the dark and stained with solution D/E. Slices were then washed, dehydrated, cleared with xylenes and cover-slipped with Permount Mounting medium (Fisher Medical, UN1294). Neurons that were identified with more complete somatic/neurite impregnation at SON and NLOT were reconstructed in a 2-D plane using a *camera lucida* additament mounted on the Nikon Eclipse 50i at 40X magnification.

### 2.7 Intracerebral cannula implantation

For implantation of permanent guide cannula into the NLOT, rats were anaesthetized with a mixture (1:1) of xylazine (20 mg/ml, Procin, Mexico) and ketamine (100 mg/ml, Inoketam, Virbac, Mexico) administered a dose of 1 ml/kg body weight intraperitoneally. Rat was placed in a stereotaxic frame (Kopf Instruments, Tujunga, CA, USA). Body temperature was maintained at 37 °C using a CMA/150 temperature controller (CMA/Microdialysis, Stockholm, Sweden). Bilateral stainless-steel cannulae of 26 gauge (C315G, Plastics One, Roanoke, VA, USA) were implanted (stereotaxic coordinates: antero-posterior -1.4 mm from bregma, medio-lateral ± 3.2 mm, and dorso-ventral 8.6 mm from the skull surface (0.6 mm was subtracted if the measurement of the micromanipulator was from the brain surface at medio-lateral 3.2 mm and antero-posterior -1.4mm). Guide cannulae were affixed on the skull with stainless steel screws and dental acrylic cement (Laboratorios Arias, México City, Mexico) and sealed with dummy cannulae (C315DC, Plastics One). Ketorolac and ceftriaxone were administered as above mentioned for three days following the surgery. Animals were housed in individual cages and allowed to recover from the surgery for one week. Beginning the 2nd-week post-surgery, the rats were handled once daily for 5 min for three consecutive days. Rats with fallen canula, signs of infection or alteration in motricity were excluded from behavioral tests.

### 2.8 Social behavior assessment and neuronal activation after intracerebral drug administration

Behavioral tests were performed on the day 11 post-surgery. On the day of the experiment, four experimental groups with rats meeting the selection criteria (*vide supra*, n=20, N=80) were formed and administered the following drugs: control (0.9% NaCl); AVP (1 ng AVP (Sigma V9879, USA)); AVP + V1a antagonist (1 ng AVP +30 ng Manning compound (V1a antagonist, Bachem, USA)) and AVP + V1b antagonist (1 ng AVP + 10 ng SSR149415 (V1b antagonist, Axon Medchem, Groningen). The doses used here were previously reported ^(9)^. The substances were injected bilaterally using two microdialysis pumps (CMA/Microdialysis, Stockholm, Sweden) in a volume of 250 nl over five minutes, and the cannulae were kept in place for 1 min after the injection to prevent backflow and to allow for diffusion as described previously ^(26)^.

Social behavior assessment using the 3CST was started 15 min after microinjections, and the time spent in the social compartment of the 3CST and the number of approaches to the rat *stranger* were quantified as mentioned above during a test period of 10 minutes. Five rats from each experimental group were randomly chosen and perfused after 60 minutes of the 3CST, and Fos immunohistochemistry was performed as described above. The number of Fos-positive nuclei in structures known to be targets of NLOT was counted within an area of 0.2 mm^2^. The rest of the rats were sacrificed with an overdose of pentobarbital and guillotined. Brains were dissected and postfixed for canula location assessment. Scores of rats whose canula tips were placed beyond 200 µm from the perimeter of the NLOT were excluded.

### 2.9 RNAscope dual ISH assays

Rats were transcardially perfused with 4% paraformaldehyde (PFA) in 0.1M PBS (PBS tablets Sigma P-4417). Brains were immediately frozen in 2-methylbutane (isopentane, Sigma-Aldrich, cat 277258, MO, USA) cooled in powdered dry ice, sectioned at a 12 μm thickness, and mounted onto *Fisher super frost slides*. Sections were dried for one hour at 60ºC, then slides were treated with 1X target retrieval reagent (from 10x stock solution provided by the supplier) in boiling water bath for 5 min. Digestion with protease-plus was made for 15 minutes. RNAscope duplex method for dual ISH hybridization with RNA probes to identify colocalization of PACAP (probe Rn-Adcyap1, cat. 400651) with V1b receptors (probe Rn-Avpr1b, cat. 443831), V1a receptors (probe Rn-Avpr1a, cat. 402531), VGLUT1 (probe Rn-Slc17a7, cat 317001), and VGLUT2 (probe Rn Slc17a6, cat. 317011) in the NLOT and hippocampus CA2, as well as the colocalization of PACAP receptor 1 (probe Rn-Adcyap1r, cat. 466981) and VGLUT1 (probe Rn-Slc17a7, cat 317001) in the insular and gustative cortex was performed. Probes were designed and provided by Advanced Cell Diagnostics, Bio-Techne (Hayward, CA, United States). Amplification and staining steps were performed following the RNAscope.2.5 HD Assay Duplex protocol for fixed frozen sections

### 2.10 Statistical analysis

Results are expressed as means ± SEM. D’Agostino and Pearson and Kolgromov-Smirnov tests were used to evaluate the normality of the data. For the behavioral tests shown in Fig. 1B (Time spent in the social compartment) and 1C (number of approaches), the results of water deprived group did not distributed normally; thus we compared them using a non-parametric Mann-Whitney test. The data of all other experiments showed a normal distribution; thus we compared them using a one-way ANOVA, followed by a Tukey’
ss multiple comparisons test to compare the effects of pharmacological micro-infusion of AVP and combined infusion with its antagonists on social behavior and number of Fos positive nuclei. Significance in all tests was set at * p<0.05, ** p< 0.01 and *** p<0.001. All statistical analyses were computed using Prism (GraphPad Software, Version 9, San Diego, CA).

**Figure 1.**
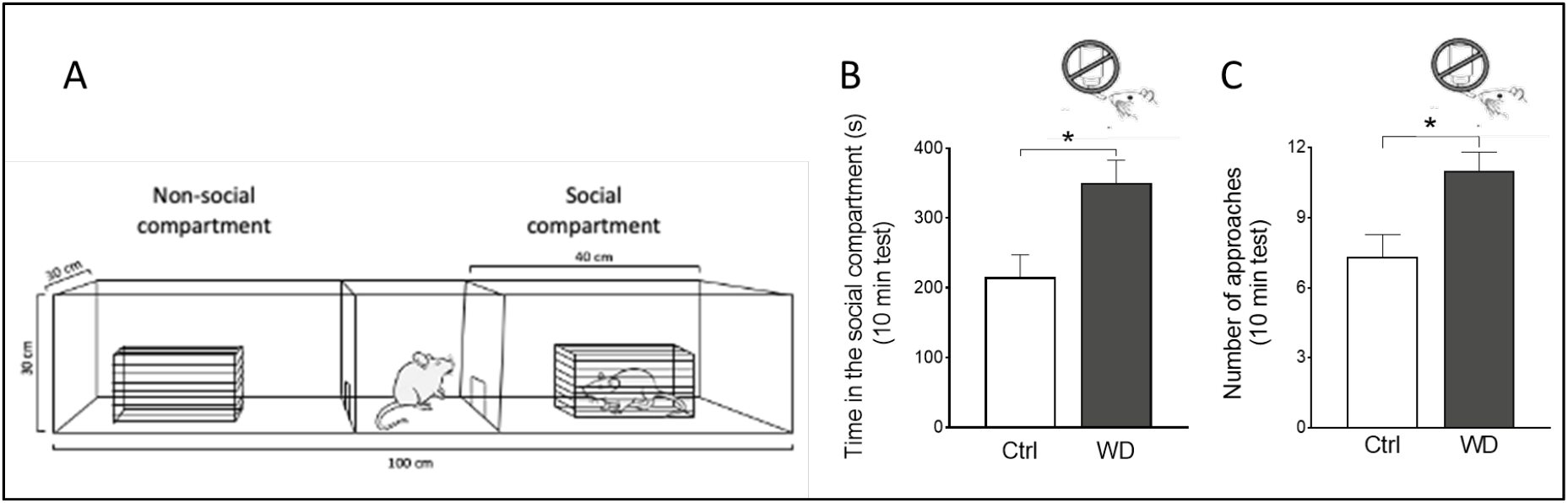
Water deprivation during 48h (WD48) increased the time of social interaction. **A:**schematic representation of the three-chamber social test (3CST) used during sociability evaluations. **B:** WD48 led to a significant increase in the time spent in the social chamber where the rat stranger was kept inside a wire cage as compared control rats. **C:** WD48 increased the number of approaches/sniffing to the rat stranger compared to the control group. Results are expressed as means ± SEM. Ctrl n= 12, WD n= 14. * p < 0.05.

## 3. RESULTS

### 3.1 WD48 led to increased time spent in the social chamber and number of approaches to the rat *stranger* during the social interaction test (3CST)

We have previously observed that WD48 increases conditional anxiety (internal, homeostatic, threat) evaluated by elevated plus maze ^(27)^ but reduces freezing behavior and increases escaping attempts when rats were exposed to a predator ^(3)^. In both cases, it seems there is a crucial involvement of the hypothalamic vasopressinergic ascending system, but in a *differential manner depending on the types of adversity the animal is facing vs. chances for individual survival*, for instance, the central amygdala pro-anxiety effect prevails in the conditional anxiety test, with main AVP sensitive neurons to be GABAergic ^(6)^ while the suppression of the lateral habenula effect prevails in the escaping of predator test ^(28)^. From these observations came the question about the social interaction consequences when the animal is under an osmotic challenge and its circuit involvement. For these reasons, we devised a simple social interaction test, the 3CST (Fig. 1A, for details see section 2.2), to evaluate if the up-regulation of the hypothalamic AVPMNN system could influence social behavior. A Mann-Whitney test showed that water-deprived rats displayed a significant enhancement in time spent in the *social* chamber where a rat *stranger* was located within a wire cage (Fig. 1B) (control: 215.5 ±31.7 vs. WD48: 350.3 ±32.4; *p*=0.013) as well as in the number of approaches/sniffing to it (control: 7.3 ±0.9 vs. 24h WD: 11 ±0.8; *p*=0.012) (Fig. 1C).

### 3.2 Fos expression triggered by WD48+3CST compared with basal, WD48 or 3CST alone depicted the social behavioral neuronal network

Fos expression analysis is a powerful instrument for getting insights into the patterns of neuronal activation during internal/external stimulations and behavioral adaptation^(29, 30)^. To investigate Fos activation, experimental subjects were submitted to 4 conditions: 1) basal condition, with food and water *ad libitum* and un-disturbed before perfusion/fixation; 2) rats underwent WD48, with food *ad libitum* (usually they stop food intake after 12h of water deprivation), un-disturbed before perfusion/fixation; 3) rats with food and water *ad libitum*, subjected to 3CST - the perfusion/fixation was performed 60min after the end of the test; 4) rats with WD48 previous to subject to the 3CST - the perfusion/fixation was performed 60min after the end of the test. Accordingly, we systematically examined increased Fos expression, as a measure of neuronal activation, throughout the brain in these four groups and made a semi-quantitative assessment (Tab. 1, see the analysis criteria in section 2.4).

As expected, social interaction caused moderate to large increases in regions involved in processing of exteroceptive and interoceptive information. These regions include the ones involved in olfactory processing such as the anterior olfactory nucleus, accessory olfactory bulb, nucleus of the lateral olfactory tract, piriform and entorhinal cortex, taenia tecta, cortical and posterior amygdala, olfactory tubercle, and endopiriform nucleus ^(31, 32)^; regions involved in visual processing such as the primary and secondary visual cortex, pretectal nucleus, the lateral geniculate complex of the thalamus and the midbrain nucleus of the posterior commissure ^(33)^; regions involved in auditory processing such as the inferior colliculus, medial geniculate nucleus and basolateral amygdala ^(34)^; limbic regions that participate in the integration of emotional states such as central amygdala, bed nucleus of the stria terminalis, accumbens, ventromedial hypothalamus and septum; regions involved in memory and coding of space information such as the hippocampal formation, entorhinal cortex and retrosplenial cortex ^(35, 36)^; and deep hypothalamic nuclei involved in stress, aggression, arousal regulation such as the paraventricular and ventromedial hypothalamic nuclei.

In contrast, activation of neurons by water deprivation was dramatically greater in SON, and PVN, two nuclei rich in osmosensitive AVPMNNs, whereas the SCN, containing non-osmosensitive vasopressinergic neurons, and other non-vasopressin-expressing nuclei of brain, exhibited far less activation in response to water deprivation (Table 1). A high number of Fos-positive nuclei was found in the BNST, a region that has been reported to be targeted by projections from osmosensitive glutamatergic neurons of the subfornical organ ^(37, 38)^.

**Table 1.**
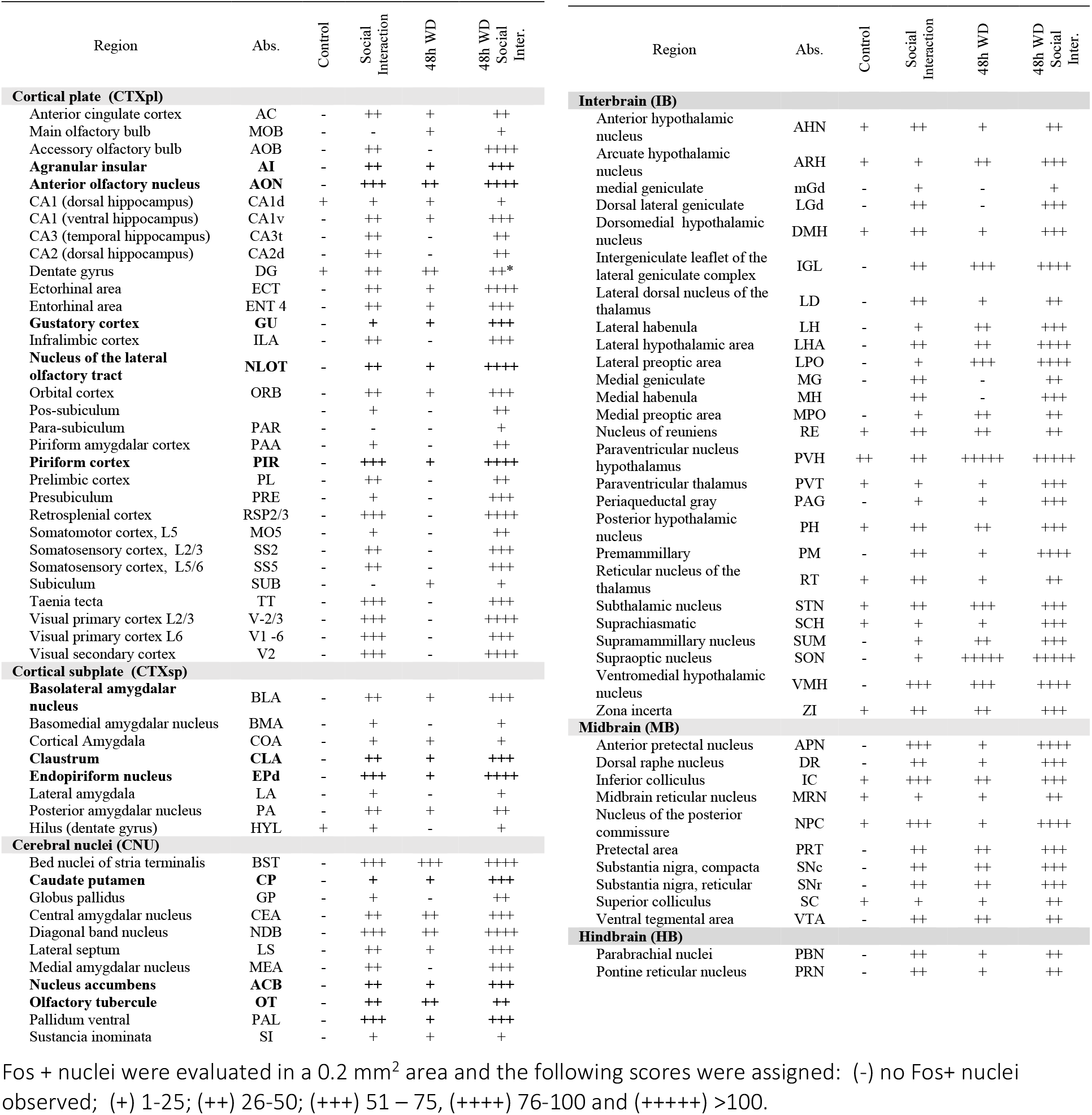
Semi-quantitative assessment of neural activation in 4 experimental conditions

WD48+3CST induced spatial neuronal activation pattern coincided with the main efferent regions of SON and the nucleus of the lateral olfactory tract (NLOT). Among these regions, the NLOT was of our particular interest, since we have noted this region to contain vasopressinergic fibers in which WD48 visibly increase IHC visibility of the AVP+ fibers (*vide infra*). The main efferent cortical regions of NLOT (Fig. 5A and supplemental figure 1) showed strong increase of Fos expression after WD48+3CST. These regions include anterior olfactory nucleus, piriform cortex, agranular insular cortex, gustative cortex, dorsal insular cortex, basolateral amygdala, claustrum, and endopiriform nucleus (see table 1, bold lettered regions).

### 3.3 Forty-eight hours of water deprivation (WD48) potentiated the hypothalamic vasopressinergic system and activate Fos in neighboring NLOT

We therefore further examined the source of origin vasopressinergic fibers as well as the nature of Fos-positive cells in NLOT after water deprivation. Concordant with progressive activation of secretion of AVP from posterior pituitary (Fig. 2A) following water deprivation, there was an increase in Fos expression both in PVN and SON (Fig. 2B’), as noted in Table 1, and in NLOT as well (Fig. 2B). Furthermore, AVP-positive terminals were found in close apposition to Fos-positive cells in NLOT (Fig. 2B‘s’), and the density of vasopressin immunopositive terminals in NLOT was correspondingly increased (Fig. 2C vs. 2D). It seems that upregulating the vasopressinergic system by osmotic stress facilitated the observation of AVP fibers in the NLOT.

**Figure 2:**
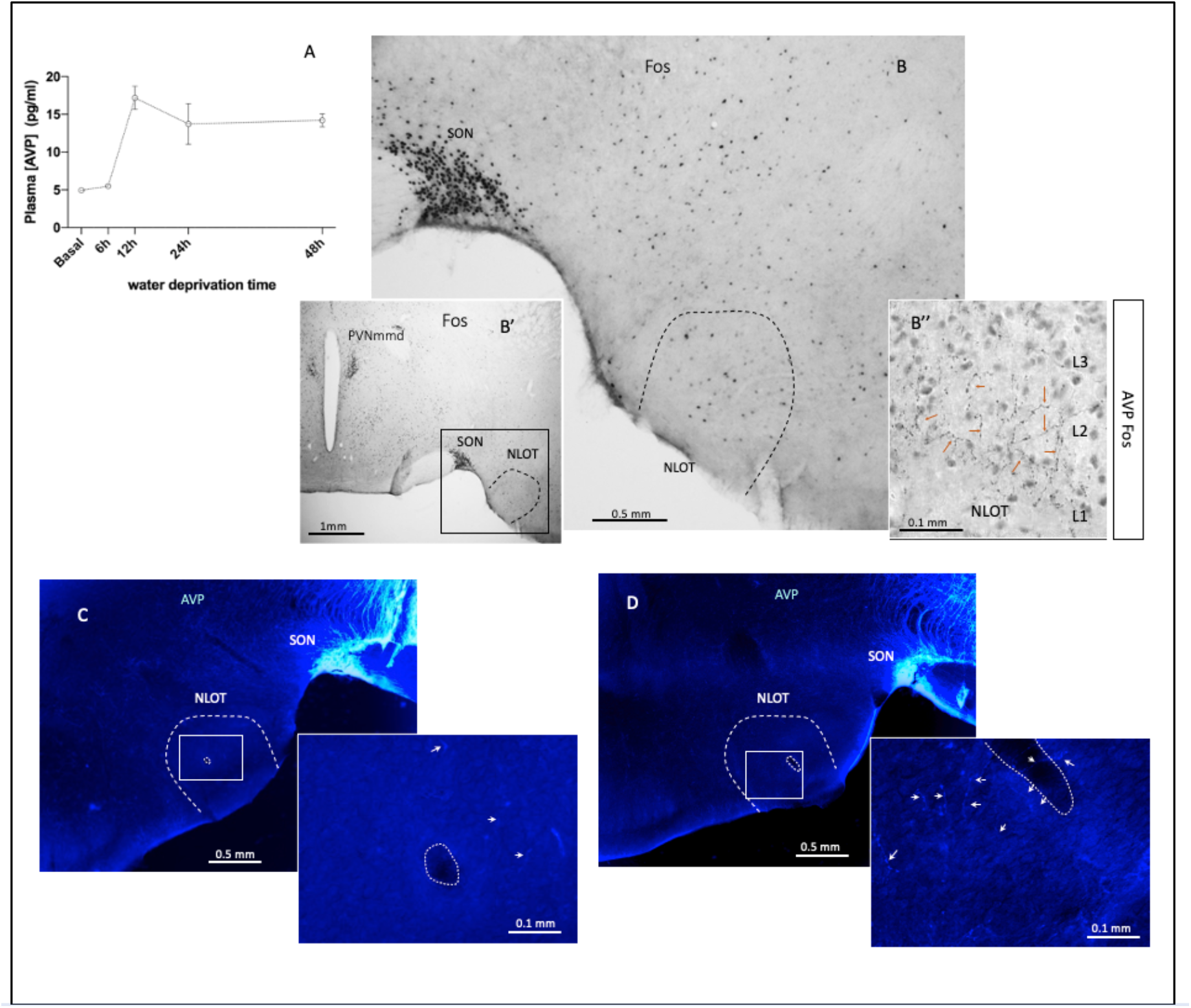
Forty-eight hours of water deprivation (WD48) potentiated the hypothalamic vasopressinergic system and activated Fos in neighboring NLOT. **A:**Time course of dynamic changes in plasma arginine vasopressin (AVP) concentration during 48 h of water deprivation, measured in wild-type male rats (n=5) with ELISA (modified from Zhang et al, 2020, with permission). Bs: example of Fos expression in hypothalamus at supraoptic level and cortical amygdalar region (B’: low magnification of B and B’’, Fos and AVP IHC in NLOT after WD48). C and D: photomicrographs showing AVP immunoreactivity (ir) (digital photos in negative mode to enhance AVP immunopositive fibers’ visibility) in the same region of B, in control and WD48 rats. Note in the high magnification insets that after WD48, the AVP + fibers clearly increased their visibility (white arrows). A vessel within the magnified region is indicated in dotted lines for anatomical reference. Abbreviatures: PVNmmd: paraventricular nucleus, medial magnocellular division; SON: supraoptic nucleus; NLOT: nucleus of lateral olfactory tract

### 3.4 NLOT received vasopressinergic input from SON of the hypothalamus revealed by Fluoro-Gold (FG) retrograde tracing and Golgi-Cox-stained sample analysis

Retrograde tracing in conjunction with staining for AVP was employed to demonstrate that SON AVPMNNs project to NLOT. Iontophoretic FG injection targeted to NLOT (see section 2.5) resulted in strong labeling in SON (Fig. 3A and B). Most of labeled cells were immunopositive to AVP (Fig. 3B). Microscopical observation of Golgi-Cox stained samples of the rat brain coronal section (150 µm of thickness), containing NLOT and the SON, with only sparse cells being stained (Fig. 3C and 3D), revealed that SON magnocellular neurons could give raise as many as 3 main axons which coursed medial, lateral and dorsally. Figure 3Da and 3Db show two magnocellular neurons, single-stained. Those two cells are called cells “a” and “b” in figure 4. Each of the cells emitted 3 main axons, numbered 1, 2, 3, which coursed medial, dorsal, and laterally. Fig. 3Dc shows the axon coming from cell “b”, axon #3, with huge varicosity, a characteristic of the magnocellular neurons, entering the layer 1 of NLOT (Fig. 4F)

**Figure 3.**
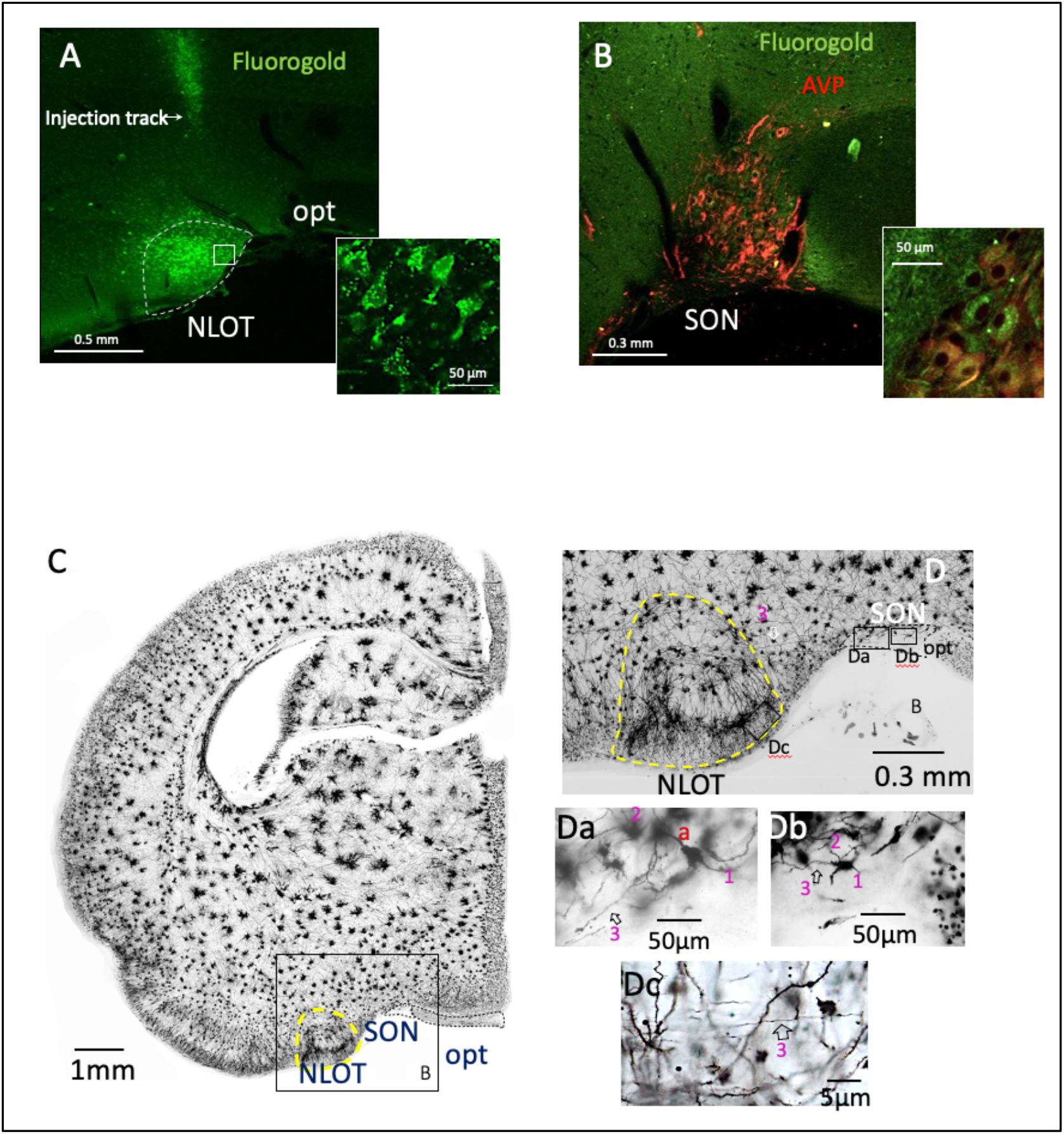
Anatomical relationship and interconnections between rat hypothalamic supraoptic nucleus (SON) and the nucleus of lateral olfactory tract (NLOT). **A** and B: retrograde tracer fluorogold injected to NLOT labeled the SON vasopressin immunoreactive neurons. C. Golgi-Cox-staining of the rat brain coronal section (150 µm of thickness) containing NLOT and the SON. D: higher magnification of C with squared region with NLOT and SON showed. Note the clearly visible allocortical feature, three layers of the NLOT clearly visible. Da and Db show two magnocellular neurons, single-stained within the SON, that they emitted 3 main axons from soma or proximal dendrites and coursed medially (1), dorsally (2) and laterally (3) that could be followed and reconstructed using adjacent sections (see Fig. 4A). Dc: showing the axon coming from cell “b”, axon #3, with huge varicosity, which is a characteristic of the magnocellular neurons, entering the layer 1 of NLOT, could be followed in the two adjacent sections. Hollow arrows indicate the main axons coming out from the magnocellular cells a or b.

**Figure 4.**
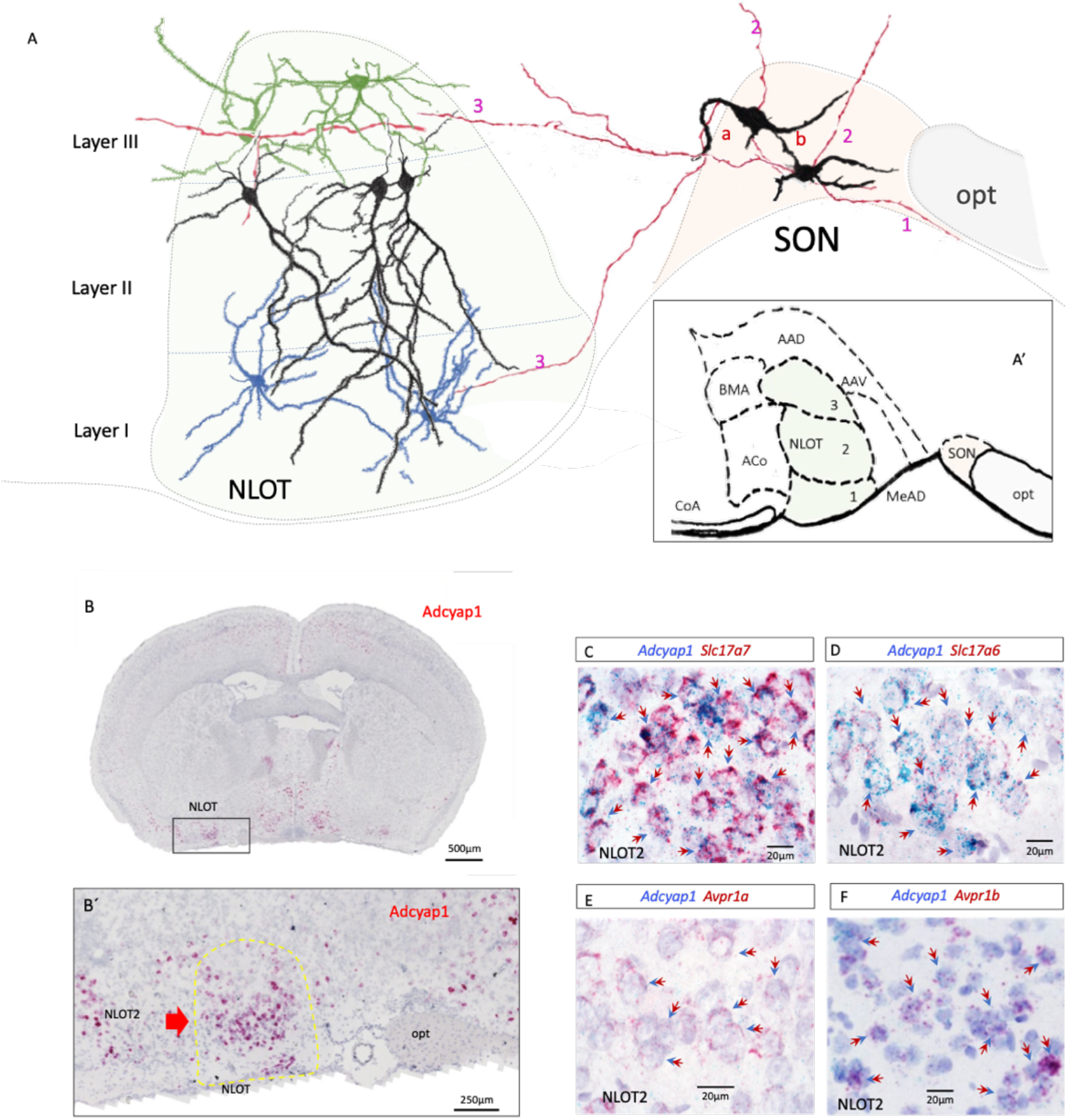
Molecular signature of NLOT principal neurons (layer 2, NLOT2), glutamatergic, PACAPergic and V1a and V1b mRNA expressing. A:camera lucida reconstructions of selective NLOT neurons, from Golgi-Cox-stained sample of Fig. 2C. Note the NLOT2 neurons (black neurons) are mainly pyramidal neurons with the apical dendrites and their subsequent branching toward the layer 1 and the brain ventral surface where most of AVP immunopositive fibers were observed. Blue cells were camera lucida-reconstructed from the NLOT1 and green cells from the NLOT3. Two SON (light beige shading magnocellular neurons with three axons emitted from cell bodies or proximal dendrites, followed in the two adjacent sections. “1” indicate the axons coursing toward the infundibulum, “2” indicate the axon coursing dorsally and “3” indicate axons coursing laterally toward NLOT (light green shading) and cortical amygdala (CoA). B: coronal section showing the RNAscope ISH for Adcyap1, mRNA encoding PACAP, at the nucleus of lateral olfactory tract (NLOT) antero-posterior level. The NLOT strongly expresses PACAP mRNA. B’: amplification of squared region in B. Note that most of PACAPergic neurons are the layer 2 neurons (NLOT2). C-F, dual in situ hybridization (DISH) using RNAscope duplex method showing the PACAPergic neurons of NLOT2 co-express Slc17a7 (mRNA for VGLUT1), Slc17a6 (mRNA for VGLUT2), Avpr1a (mRNA for vasopressin receptor V1a) and Avpr1b (mRNA for vasopressin receptor V1b). Doble arrows indicate examples of co-expression.

### 3.5 Molecular signature of NLOT principal neurons (layer 2, NLOT2) and projection fields reveals its cooperative role with hypothalamic vasopressinergic signaling and NLOT PACAPergic signaling pathway

The NLOT is an isolated tri-laminar ovoid cell mass located in the anterior cortical amygdalar region ^(19, 39)^. Nissl staining subdivides the NLOT nucleus in three cell layers. Layer 1 (NLOT1) is a subpial molecular zone with scattered neurons, which receives mitral cell input from the main olfactory bulb. Layer 2 (NLOT2) is a thick and dense corticoid aggregate of pyramidal neurons of medium size with apical dendrites entering NLOT1 (Fig. 4F, for example). Layer 3 (NLOT3) is a multiform layer disposed deep to layer 2. It contains a mixture of small and large neurons, some of them possibly representing inhibitory interneurons of subpallial origin ^(21)^. We next explored the neurochemical signature(s) of the neurons in NLOT that are the presumptive targets of vasopressinergic terminals from SON, which terminate mainly in the layer 1 where the pyramidal neurons apical dendrites branch. Figure 4A shows three types of neurons’ morphology reconstructed using camera lucida and Golgi-Cox stained samples (section thickness: 150 µm). Pyramidal neurons of the NLOT2 are of interest to this study. Using dual ISH (RNAscope duplex method), we identified a population of PACAPergic, VGLUT1 and VGLUT2 expressing neurons in layer II of NLOT (Fig. 4C and 4D) ^(40)^ representing pyramidal neurons (black cells) which, upon Golgi staining and reconstruction, send dendritic processes into both layers I and layer III of NLOT where the SON magnocells send their axons (Fig. 4A, red axons reconstructed with camera lucida in 3 adjacent sections of thickness 150 µm each). As these neurons also express the vasopressin receptors V1a and V1b (Fig. 4E and 4F), they are candidates as the target neurons of the vasopressinergic projections from SON. In fact, the Fos positive neurons in the layer II are likely among those PACAPergic (and glutamatergic) neurons after 48WD.

Cortical efferent regions of NLOT2 PACAPergic pyramidal neurons which also express VGLUT2 ^(40)^ (Fig. 4D) and Sim1 gene ^(41)^ are revealed by Allen connectivity atlas (Fig. 5A and supplementary Fig. 1). Figure 5A is a map of NLOT projections made with a mouse line <u>Adcyap1-2A-Cre, Exp. 187269162</u>. The main efferent regions include anterior olfactory nucleus, piriform cortex, agranular insular cortex, gustative cortex, dorsal insular cortex, basolateral amygdala, claustrum, and endopiriform nucleus (see table 1, bold lettered regions) which all strongly express the PACAP receptor PAC1 mRNA (Adcyap1r1, Fig. 5B) and also co-express VGLUT1 mRNA (Slc17a7, Fig. 5C and insets).

**Figure 5.**
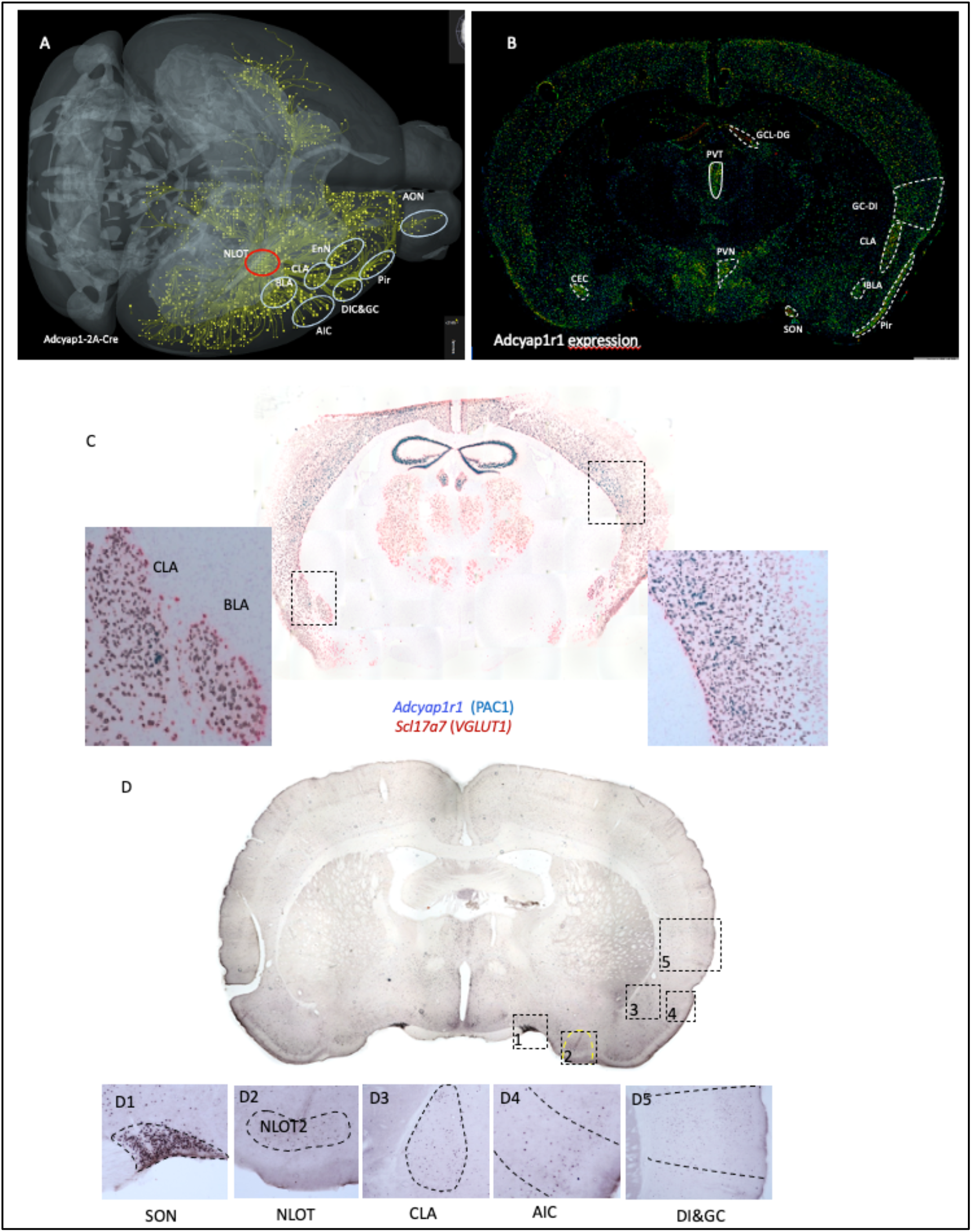
Main efferent regions of glutamate-PACAPergic NLOT analyzed in this study strongly co-express mRNAs of PACAP receptor PAC1 (Adcyapr1r) and VGLUT1 (Slc17a7) and were activated by 48WD. A:Axonal projections from PACAP transfected NLOT neurons. Obtained from Allen mouse brain connectivity atlas (Experiment: 187269162 Transgenic line: Adcyap1-2A-Cre). B: coronal section showing the expression of Adcyap1r1 (PAC1 receptor mRNA) with cortical regions that are target of NLOT delineated in dotted lines. Image obtained from Allen ISH brain atlas (Experiment: 74988667). C: Coronal section processed for dual ISH using Duplex RNAscope, to show the colocalization of VGLUT1 and PAC1 in cortical regions targeted by projections from PACAPergic NLOT neurons. D: Up-regulation of Fos in putative projection targets of NLOT PACAPergic neurons after WD48+3CSI. Panels: D1 – D5 show the key regions discussed in this study: SON: supraoptic nucleus; NLOT: nucleus of lateral olfactory tract; CLA: claustrum; AIC: Agranular insular cortex; DI, dorsal insular cortex; GC: gustative cortex. The whole brain Fos expression semi-quantitative analysis is reported in table 1.

### 3.6 Increased social behavior and Fos expression induced by vasopressin micro-infusion in NLOT is blocked by both V1a and V1b receptor antagonists

In order to ascertain whether or not AVP can directly modulate the activity in NLOT and influence social behavior, AVP (1 ng/side) was bilaterally infused into this region (Fig. 6A) 15 min before 3CST. To evaluate the participation of the V1b and V1a receptors, the pharmacological antagonists SSR149415 (V1b antagonist) and Manning compound (V1a antagonist) were co-injected with vasopressin at 30 ng/side in 20 rats per condition. Histological verifications were made for sample inclusion assessment. As a histological inclusion criterion for Fos expression and behavioral test analysis, the cannula tips were confirmed to be located within 200 µm perimeter of NLOT boundaries.

**Figure 6.**
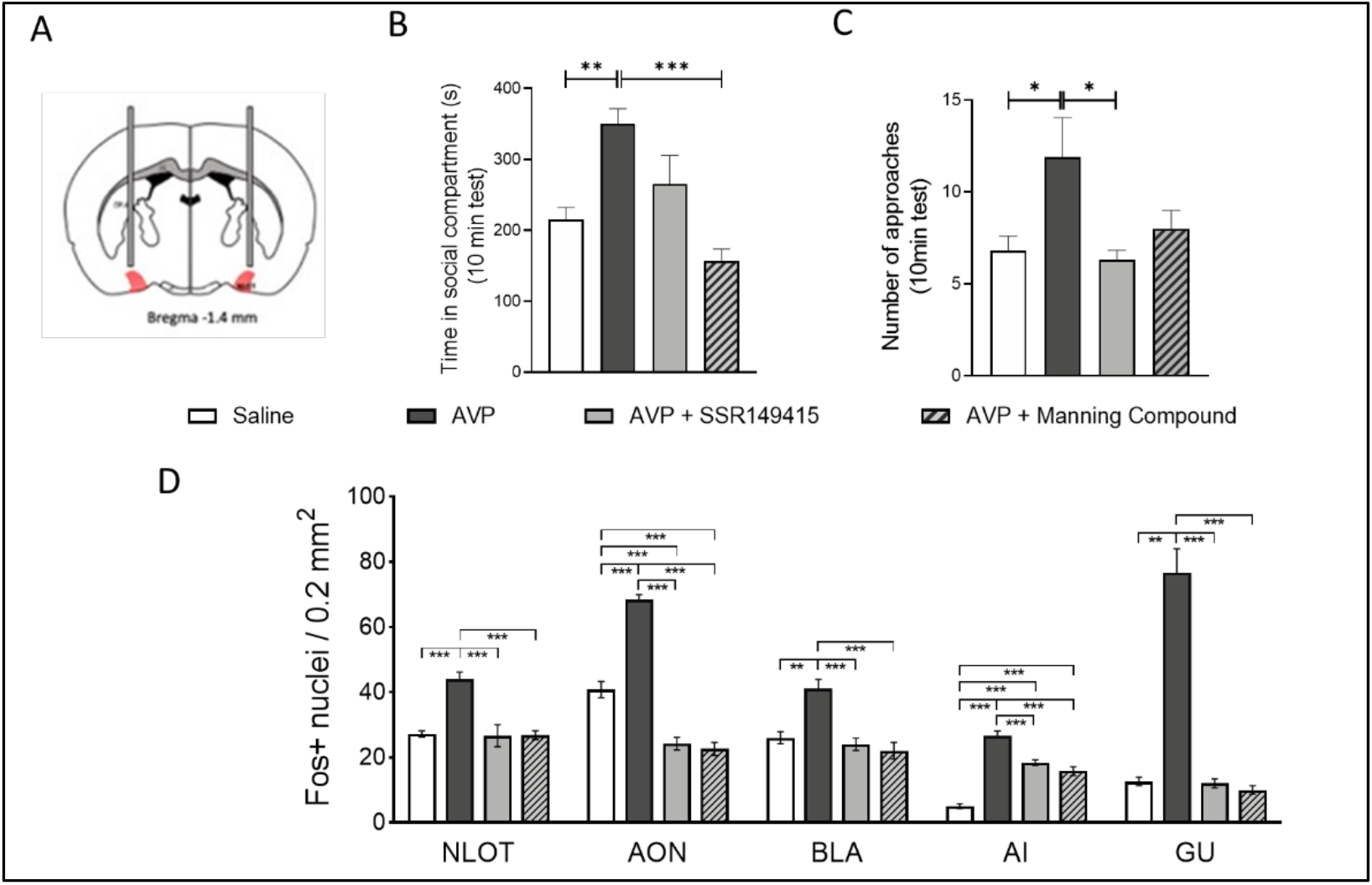
Vasopressin microinfusion into the nucleus of the lateral olfactory tract (NLOT) induces both an increase in social behavior and Fos+ nuclei expression. **(A)** Representative scheme showing a bilateral cannula placeman within the NLOT. **(B)** AVP infusion (1ng/side) increased the time that the rats spent inside the social compartment these effects were fully prevented by the simultaneous infusion of both AVP V1b receptor (10 ng/side, SSR149415 (SSR) antagonist, and AVP V1a receptor (10 ng/side, Manning compound antagonist. **(C)** A significant difference was found in the number of entries between the infusion of AVP vs the saline group these effects were blocked by the simultaneous infusion of AVP V1b and V1a receptor antagonist. **(D)** AVP infusion into NLOT elicited an enhancement of Fos+ expression in AON, BLA, AI, and GU, as compared with the respective saline-treated group. The simultaneous infusion of AVP+V1b and AVP+V1a antagonist resulted in a decreased Fos expression in all the areas studied in this work. Abbs: NLOT: nucleus of lateral olfactory tract; AON: accessory olfactory nucleus; BLA: Basolateral amygdala; AI: agranular insular; GU: gustatory area. Results are expressed as means ± SEM. **P< 0.01; ***P<0.001. Saline n=20; AVP n=19; AVP+SSR n=19; AVP+Manning C. n=19. Fos expression quantification (n= 5 rats for each group).

A one-way ANOVA showed a significant effect of the drug infusion on the time spent in the social compartment (Fig. 6B) (F (3,74) = 10, *p*<0.0001). A post-hoc Tukey’
ss test showed that AVP significatively increased the time (in sec) that the experimental rats spent in the *social* compartment as compared with the saline-treated group (AVP: 350.3 ±21.3 *vs*. control: 215.4 ±16.9, *p*<0.01). This effect was abolished when vasopressin was co-administered with SSR149415 (265.3 ±40.3) or Manning compound (156.3 ± 17.6). With respect to the number of approaches, the drugs infusion produced significant effects on the number of approaches to the rat in the social compartment (Fig. 6C) (F3,74) = 3.8, *p*=0.012). Tukey’s post-hoc test uncovered a significant increase in this approach behavior posterior to applying AVP compared with saline group (AVP: 11.9 ±2.2 *vs*. 6.8 ±0.8; *p*<0.05). The co-infusion of AVP+ SSR149415 prevented the increase in the number of approaches to the social compartment (6.3 ±0.5). Finally, the microinfusion of Manning compound decreased the number of approaches (8 ±1) induced by AVP.

To confirm that NLOT PACAPergic/glutamatergic neurons are plausible targets for activation by vasopressin projections from SON, we examined Fos elevation in selected regions known to be involved in social behavior (Fig 6D). One-way ANOVA showed a significant effect of treatment in all evaluated regions (Fig. 6D): NLOT: F(3,16) = 15, *p*<0.0001; AON: F(3,16) = 111.6, *p*<0.0001; BLA: F(3,16) = 13.6, *p*=0.0001; AI: F(3,16) = 60.4, *p*<0.0001; GU: F(3,16) = 69.9, *p*<0.0001. Post-hoc Tukey multiple comparison test showed that bilateral AVP (1 ng/side) infusion into the nucleus of the lateral olfactory tract (NLOT) elicited an increase of Fos+ expression in NLOT (saline: 27.2 ±1, vs AVP: 44 ±2.2, *p*<0.001), no significant differences were observed between saline and AVP+SSR149415 (26.6 ±3.4) nor between saline vs. AVP + Manning compound (26.8 ±1.4). In the anterior olfactory nucleus (AON) we found that compared to saline (40.8 ±2.5), AVP increased the number of Fos positive nuclei (68.4 ±1.5, p<0.0001), and the application of AVP with either of the antagonists produced a significant decrease in the number of Fos+ nuclei (AVP+SSR149415: 24.2 ±1.9 *vs* saline: 40.8 ±2.5, *p*<0.001) and (AVP + Manning compound: 22.6 ±2.0 *vs*. saline: 40.8 ±2.5, *p*<0.0001). In the case of the basolateral amygdala (BLA), AVP elicited significant increases in Fos activation (saline: 26 ±1.9 *vs. AVP:* 41 ±2.9, *p*<0.01) and this effect was blocked with AVP+SSR149415 (24 ±1.9) and with Manning compound (22 ±2.6). For agranular insular cortex (AI), AVP increased the number of Fos+ nuclei (saline: 5 ±0.7 *vs*. AVP: 26.6 ±1.5; p<0.0001); however AVP antagonists did not blocked this increase (AVP+SSR149415: 18.4 ±0.8 *vs*. saline: 5 ±0.7, p<0.0001) and (AVP + Manning compound: 15.8 ±1.4 *vs*. saline: 5 ±0.7, p<0.0001). In gustatory area (GU), AVP significantly increased the number of Fos+ nuclei (Saline: 12.6 ±1.3 *vs. AVP:* 76.6 ±7.4, p<0.0001) and V1a and V1b antagonists abolished this effect as no differences were found between saline and SSR14915 (12 ±1.4), or Manning compound (10 ±1.3).

## DISCUSSION

In this study, we focused on a newly-characterized projection system from AVPMNNs of the supraoptic nucleus (SON) of the hypothalamus to the nucleus of the lateral olfactory tract (NLOT), the activation of these neurons by water deprivation, and ensuing effects on their neurochemistry, their connections, and their potential effects on social behavior.

We report here that rats stressed osmotically via WD48 had a higher score in sociability than euhydrated rats. Osmotic stress potently increases the metabolic activity of AVPMNNs in the hypothalamus, and AVP mRNA expression in PVN and SON correlates with changes in social interaction in mice ^(42, 43)^. Intracerebroventricular infusion of AVP increases social contact in prairie voles ^(44)^ and novel conspecific social interaction in the Syrian hamster ^(45)^. Brattleboro rats lacking AVP due to a genetic prohormone processing defect, show various behavioral impairments, including altered social development ^(14)^ and less social playing and prosocial ultrasonic vocalizations ^(13)^. The duration of social recognition in rats ^(46)^ is dependent on the levels of AVP in the LS. Variation in expression of vasopressin receptor genes (AVPR1A, AVPR1B) correlates with variation in social behavior in rhesus macaques ^(47)^. Here, we identify for the first time SON AVPMNN direct innervation of the NLOT, revealing a candidate circuit activated by osmotic challenge induced by water deprivation that is capable of conveying information to downstream brain circuitry involved in social behavior. We also observed increased Fos expression in the nucleus of the lateral olfactory tract (NLOT) and its efferent cortical regions after WD48 + 3CST, indicating that the homeostatic state is able to modulate neocortical social interaction processing centers via NLOT, a hub of cortical amygdalar complex. We confirmed this result is AVP-mediated by injecting vasopressin directly into the NLOT. A novel finding in this study was that the V1a and V1b receptor antagonists co-treatment reduced the AVP-induced increase in sociability. The fact that this phenomenon was blocked by either V1a or V1b receptor antagonists co-infused with AVP, suggests that both types of receptor are necessary, while either one alone is insufficient, to mediate these behavioral effects.

NLOT projections to the accessory olfactory system which are activated during the social investigation behavior in which AVP participates through neuronal signaling to filter social cues of odor ^(48)^. Similarly, NLOT-lesioned rats spend significantly less time sniffing out odors associated with social contacts than intact rats regardless of the number of times exposed to the olfactory stimulus ^(49)^. It is known that water deprivation enhances both AVP-immunoreactivity and Fos expression in the hypothalamic paraventricular and supraoptic nuclei ^(5)^ and therefore vasopressin effects could be exerted elsewhere than at the level of the amygdala. However, the fact of that exogenous administration of AVP into NLOT could produce identical behavioral effects as that induced by WD, support the conclusion that the effects of WD are mediated by AVP release in the NLOT. The effects of AVP microinfusion on Fos activation in AON, BLA, AI, and GU, and blockade by AVP antagonist co-administration in NLOT alone, is also consistent with the notion that AVP affects activation of this behavioral network by its action as a neurotransmitter at the NLOT ‘hub’. A previous study documented enhancement of social interaction after osmotic dehydration (subcutaneous injection of 2 M NaCl), and it seems likely that this osmotic stressor, like water deprivation, causes these effects on social behavior also through AVP release in NLOT ^(50)^.

It is uncertain at the present time how altered social behavior (increased social interaction) after water deprivation represents an integration of homeostatic and allostatic drive that is beneficial to animal survival (and would, at least simplistically) explain the evolution and stabilization of the pathway described here in rat. Broadly, increased attention to the behavior and proximity of con-specifics during resource scarcity may have both positive and negative effects. For example the borer and social mutations of another neuropeptide-liganded GPCR, the NPY-like receptor of C. elegans, are posited to drive foraging behavior when food is scarce, and social behavior (and increased mating) when food is plentiful, respectively ^(51)^. In this context, increased sociability during water deprivation would seem to be disadvantageous, rather than the reverse. However, it must be borne in mind that the behavioral model chosen here as a ‘behavioral readout’ for water deprivation (social interaction) may be insufficiently rich/complex to capture the full range of advantages and disadvantages of increased social interaction during periods of scarcity, or indeed if ‘water deprivation’ is in fact a model for anything resembling resource scarcity. For example, relative preferences for brackish versus non-brackish water seem to be a behavioral characteristic that involves vasopressinergic neurons of the hypothalamus, in particular a dynorphinergic component of vasopressin neurotransmission in hypothalamus^(2)^. Thus, we postpone speculation on the relative survival advantages of a linkage between social behavior and water deprivation, and note here only that this linkage exists, at least in part, because of a vasopressinergic projection from SON to NLOT, in the rat.

Besides the vasopressinergic SON**→**NLOT PACAPergic projection system described here, there are at least two other AVPMNN systems that link homeostasis and behavior that have been recently reported ^(3, 6, 28, 52)^. In these latter cases, the AVP target neurons (GERNs in LHb, amygdalar neurons, noradrenergic neurons) have their own well-established downstream connections that suggest how vasopressinergic inputs to them ultimately affect behavior. For the present case, it is much less clear, because the downstream connections of NLOT2 PACAP/glutamate neurons are not well-established.

It is worth noting that receptor-specific pharmacological manipulation of this pathway (and parallel ones) for potential therapeutic purposes could be easily separate vasopressin-like effects on osmoregulation itself (mediated mainly via V2), from behavioral effects. It will be of interest in future experiments to determine whether or not a combination of V1a and V1b *agonists* can mimic the behavioral effects of AVP infusion into the brain on behavioral independently of osmoregulatory effects intrinsic to arginine vasopressin.

Finally, we wish to note some of the limitations of our study that will require fuller exploration in the future. First, we have inferred the downstream projections of NLOT PACAPergic neurons based on their demonstration in the mouse, and because these regions demonstrate PAC1 receptor expression, and fos up-regulation after 48WD consistent with activation via an SON^AVP^**→**NLOT^PACAP^**→**neocortical regions^PAC1^. We speculate that it is this circuit that broadcasts the hydromineral status of the animal to centers mediating behavior. In fact causality will be established only when it is possible to show that activation of PACAPergic NLOT neurons is sufficient to drive the behavioral phenotype elicited by water deprivation, and that activation of these neurons is driven exclusively via SON vasopressinergic projections to NLOT. It will also be of interest to define more clearly the mechanism whereby a combined activation of V1a and V1b receptors is required for elicitation of NLOT-mediated alteration in social behavior

## Acknowledgments

We acknowledge support from DGAPA-UNAM (PAPIIT-IN216918, IG200121, LZ) and CONACYT (CB-238744) to LZ. MH002386, NIMH, NIH, USA (LEE). OH-P and MAZ were supported by Post-Doctoral Scholarship Program at UNAM (POSDOC-DGAPA).

## Supplementary information

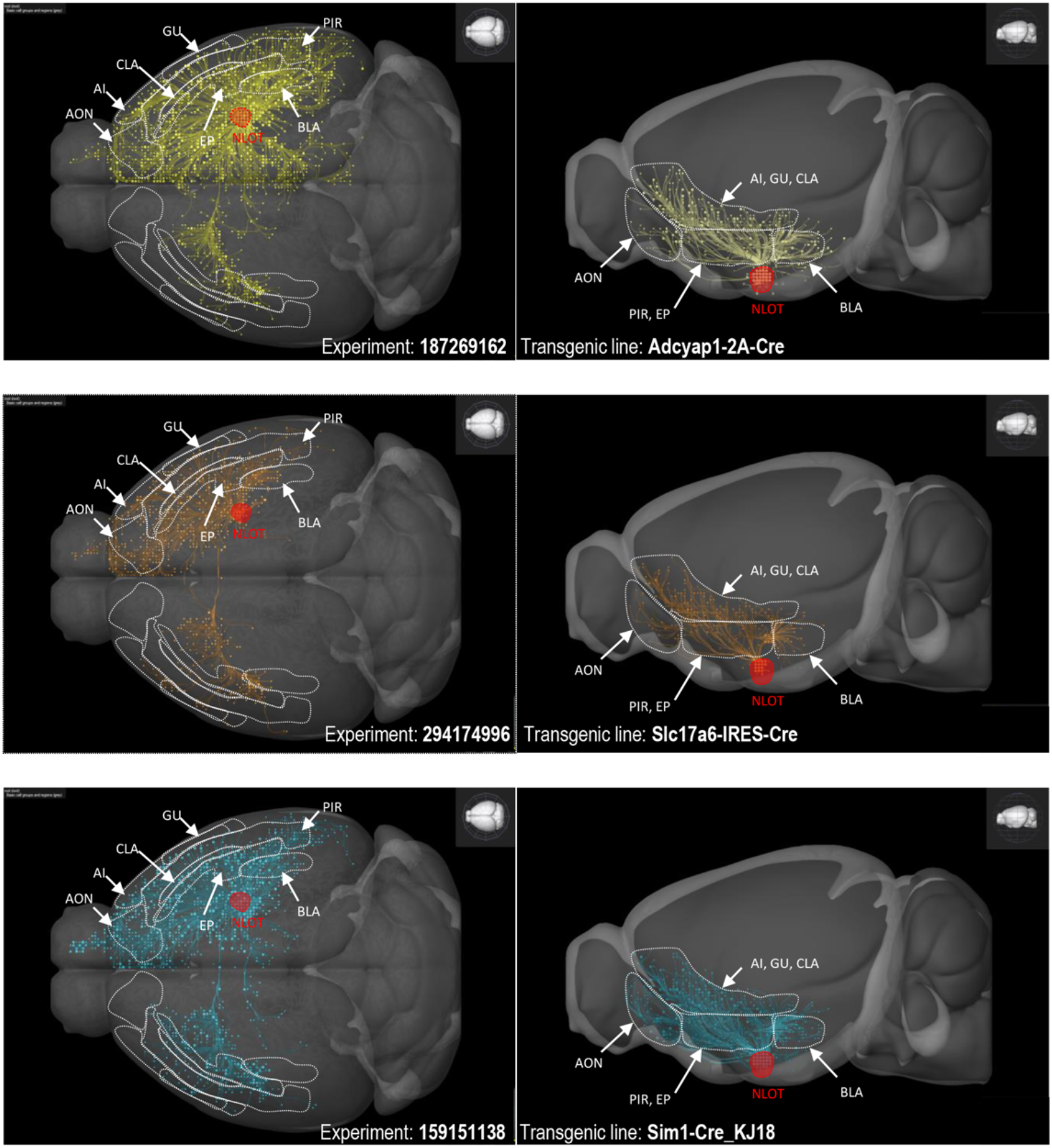
Nucleus of lateral olfactory tract (NLOT) neurons with different neurochemical markers share similar projection targets in cortical regions. Three dimensional projection patterns in dorso-ventral (left) and sagittal (right) views of NLOT virally transfected neurons in which reporter gene expression was restricted by selective CRE expression driven by the promoters of PACAP (Adcyap1 2A Cre) shown here in yellow, VGLUT2 (Slc17a6 IRES Cre) shown in orange or *single minded homolog 1* (Sim1 Cre_KJ18) depicted in blue. Notice that for all the experiments the cortical regions that were presented in bold letters in the table 1 (delineated by white dotted lines) received strong innervation by glutamatergic / PACAPergic axons originated in NLOT (delineated by red dotted lines). All of these target regions displayed increased Fos activity after 48WD+3CST. Reconstructions of the projection pathways were generated using the *Brain Explorer 2* program of the Allen Institute for Brain Science and the data of the experiments 187269162, 294174996 and 159151138 that can be found in the *Mouse Brain Connectivity Atlas*. Abbreviations: AON: anterior olfactory nucleus, AI: agranular insular cortex, BL A: basolateral amygdala, CLA: claustrum, GU: gustatory cortex, NLOT: nucleus of the lateral olfactory tract, PIR: piriform cortex.

## Notes

Supported by grants: UNAM-DGAPA-PAPIIT- PAPIIT-IN216918 & GI200121 & CONACYT-CB- 283279 (LZ); MH002386, NIMH, NIH, USA (LEE).

### Competing Interest Statement

The authors have declared no competing interest.

### Summary of Updates

Statistics amendments and discussion on limitations of this study.

## REFERENCES

1. Eiden LE, Hernández VS, Jiang S, Zhang L. Neuropeptides and Small-molecule Amine Transmitters: Cooperative Signaling in the Nervous System. Cellular and Molecular Life Sciences. 2022, in press.

2. Zhang L, Hernández VS, Murphy D, Young WS, Eiden LE. Fine chemo-anatomy of hypothalamic magnocellular vasopressinergic system with an emphasis on ascending connections for behavioural adaptation. In: Valery Grinevich ÁD, ed. Neuroanatomy of Neuroendocrine Systems Springer-Nature 2021.

3. Zhang L, Hernandez VS, Vazquez-Juarez E, Chay FK, Barrio RA. Thirst Is Associated with Suppression of Habenula Output and Active Stress Coping: Is there a Role for a Non-canonical Vasopressin-Glutamate Pathway? Front Neural Circuits. 2016; 1013.

4. Zhang L, Padilla-Flores T, Hernandez VS, Zetter MA, Campos-Lira E, Escobar LI, Millar RP, Eiden LE. Vasopressin acts as a synapse organizer in limbic regions by boosting PSD95 and GluA1 expression. J Neuroendocrinol. 2022e13164.

5. Zhang L, Medina MP, Hernandez VS, Estrada FS, Vega-Gonzalez A. Vasopressinergic network abnormalities potentiate conditioned anxious state of rats subjected to maternal hyperthyroidism. Neuroscience. 2010; 168(2): 416–28.

6. Hernandez VS, Hernandez OR, Perez de la Mora M, Gomora MJ, Fuxe K, Eiden LE, Zhang L. Hypothalamic Vasopressinergic Projections Innervate Central Amygdala GABAergic Neurons: Implications for Anxiety and Stress Coping. Front Neural Circuits. 2016; 1092.

7. Hernandez VS, Vazquez-Juarez E, Marquez MM, Jauregui-Huerta F, Barrio RA, Zhang L. Extra-neurohypophyseal axonal projections from individual vasopressin-containing magnocellular neurons in rat hypothalamus. Front Neuroanat. 2015; 9130.

8. Campos-Lira E, Kelly L, Seifi M, Jackson T, Giesecke T, Mutig K, Koshimizu TA, Hernandez VS, Zhang L, Swinny JD. Dynamic Modulation of Mouse Locus Coeruleus Neurons by Vasopressin 1a and 1b Receptors. Front Neurosci. 2018; 12919.

9. Hernandez-Perez OR, Crespo-Ramirez M, Cuza-Ferrer Y, Anias-Calderon J, Zhang L, Roldan-Roldan G, Aguilar-Roblero R, Borroto-Escuela DO, Fuxe K, Perez de la Mora M. Differential activation of arginine-vasopressin receptor subtypes in the amygdaloid modulation of anxiety in the rat by arginine-vasopressin. Psychopharmacology (Berl). 2018; 235(4): 1015–27.

10. DeVito LM, Konigsberg R, Lykken C, Sauvage M, Young WS, 3rd, Eichenbaum H. Vasopressin 1b receptor knock-out impairs memory for temporal order. J Neurosci. 2009; 29(9): 2676–83.

11. Cilz NI, Cymerblit-Sabba A, Young WS. Oxytocin and vasopressin in the rodent hippocampus. Genes Brain Behav. 2019; 18(1): e12535.

12. Caldwell HK, Wersinger SR, Young WS, 3rd. The role of the vasopressin 1b receptor in aggression and other social behaviours. Progress in Brain Research. 2008; 17065–72.

13. Paul MJ, Peters NV, Holder MK, Kim AM, Whylings J, Terranova JI, de Vries GJ. Atypical Social Development in Vasopressin-Deficient Brattleboro Rats. eNeuro. 2016; 3(2).

14. Schatz KC, Kyne RF, Parmeter SL, Paul MJ. Investigation of social, affective, and locomotor behavior of adolescent Brattleboro rats reveals a link between vasopressin’s actions on arousal and social behavior. Horm Behav. 2018; 1061–9.

15. Frank E, Landgraf R. The vasopressin system--from antidiuresis to psychopathology. Eur J Pharmacol. 2008; 583(2-3): 226–42.

16. Young LJ. Oxytocin and vasopressin as candidate genes for psychiatric disorders: lessons from animal models. Am J Med Genet. 2001; 105(1): 53–4.

17. Inyushkin AN, Orlans HO, Dyball RE. Secretory cells of the supraoptic nucleus have central as well as neurohypophysial projections. J Anat. 2009; 215(4): 425–34.

18. Zhang L, Hernandez VS. Synaptic innervation to rat hippocampus by vasopressin-immuno-positive fibres from the hypothalamic supraoptic and paraventricular nuclei. Neuroscience. 2013; 228139–62.

19. Santiago AC, Shammah-Lagnado SJ. Efferent connections of the nucleus of the lateral olfactory tract in the rat. J Comp Neurol. 2004; 471(3): 314–32.

20. McDonald AJ. Cytoarchitecture of the nucleus of the lateral olfactory tract: a Golgi study in the rat. Brain Res Bull. 1983; 10(4): 497–503.

21. Garcia-Lopez M, Abellan A, Legaz I, Rubenstein JL, Puelles L, Medina L. Histogenetic compartments of the mouse centromedial and extended amygdala based on gene expression patterns during development. J Comp Neurol. 2008; 506(1): 46–74.

22. Moy SS, Nadler JJ, Perez A, Barbaro RP, Johns JM, Magnuson TR, Piven J, Crawley JN. Sociability and preference for social novelty in five inbred strains: an approach to assess autistic-like behavior in mice. Genes Brain Behav. 2004; 3(5): 287–302.

23. Wee BE, Francis TJ, Lee CY, Lee JM, Dohanich GP. Mate preference and avoidance in female rats following treatment with scopolamine. Physiol Behav. 1995; 58(1): 97–100.

24. Dunn FL, Brennan TJ, Nelson AE, Robertson GL. The role of blood osmolality and volume in regulating vasopressin secretion in the rat. J Clin Invest. 1973; 52(12): 3212–9.

25. Paxinos G, and Watson, C.. The Rat Brain in Stereotaxic Coordinates Amsterdam., 2006.

26. Perez de la Mora M, Lara-Garcia D, Jacobsen KX, Vazquez-Garcia M, Crespo-Ramirez M, Flores-Gracia C, Escamilla-Marvan E, Fuxe K. Anxiolytic-like effects of the selective metabotropic glutamate receptor 5 antagonist MPEP after its intra-amygdaloid microinjection in three different non-conditioned rat models of anxiety. Eur J Neurosci. 2006; 23(10): 2749–59.

27. Zhang L, Hernandez VS, Liu B, Medina MP, Nava-Kopp AT, Irles C, Morales M. Hypothalamic vasopressin system regulation by maternal separation: its impact on anxiety in rats. Neuroscience. 2012; 215135–48.

28. Zhang L, Hernandez VS, Swinny JD, Verma AK, Giesecke T, Emery AC, Mutig K, Garcia-Segura LM, Eiden LE. A GABAergic cell type in the lateral habenula links hypothalamic homeostatic and midbrain motivation circuits with sex steroid signaling. Transl Psychiatry. 2018; 8(1): 50.

29. Dragunow M, Faull R. The use of c-fos as a metabolic marker in neuronal pathway tracing. J Neurosci Methods. 1989; 29(3): 261–5.

30. Kovacs KJ. Measurement of immediate-early gene activation-c-fos and beyond. J Neuroendocrinol. 2008; 20(6): 665–72.

31. Mori K, Sakano H. Olfactory Circuitry and Behavioral Decisions. Annu Rev Physiol. 2021; 83231–56.

32. Sugai T, Yamamoto R, Yoshimura H, Kato N. Multimodal cross-talk of olfactory and gustatory information in the endopiriform nucleus in rats. Chem Senses. 2012; 37(8): 681–8.

33. Seabrook TA, Burbridge TJ, Crair MC, Huberman AD. Architecture, Function, and Assembly of the Mouse Visual System. Annu Rev Neurosci. 2017; 40499–538.

34. Knipper M, Van Dijk P, Nunes I, Ruttiger L, Zimmermann U. Advances in the neurobiology of hearing disorders: recent developments regarding the basis of tinnitus and hyperacusis. Prog Neurobiol. 2013; 11117–33.

35. Knierim JJ. The hippocampus. Curr Biol. 2015; 25(23): R1116–21.

36. Vann SD, Aggleton JP, Maguire EA. What does the retrosplenial cortex do? Nat Rev Neurosci. 2009; 10(11): 792–802.

37. Matsuda T, Hiyama TY, Niimura F, Matsusaka T, Fukamizu A, Kobayashi K, Kobayashi K, Noda M. Distinct neural mechanisms for the control of thirst and salt appetite in the subfornical organ. Nat Neurosci. 2017; 20(2): 230–41.

38. Zimmerman CA, Leib DE, Knight ZA. Neural circuits underlying thirst and fluid homeostasis. Nat Rev Neurosci. 2017; 18(8): 459–69.

39. Krettek JE, Price JL. A description of the amygdaloid complex in the rat and cat with observations on intra-amygdaloid axonal connections. J Comp Neurol. 1978; 178(2): 255–80.

40. Zhang L, Hernandez VS, Gerfen CR, Jiang SZ, Zavala L, Barrio RA, Eiden LE. Behavioral role of PACAP signaling reflects its selective distribution in glutamatergic and GABAergic neuronal subpopulations. Elife. 2021; 10.

41. Garcia-Calero E, Lopez-Gonzalez L, Martinez-de-la-Torre M, Fan CM, Puelles L. Sim1-expressing cells illuminate the origin and course of migration of the nucleus of the lateral olfactory tract in the mouse amygdala. Brain Struct Funct. 2021; 226(2): 519–62.

42. Murakami G, Hunter RG, Fontaine C, Ribeiro A, Pfaff D. Relationships among estrogen receptor, oxytocin and vasopressin gene expression and social interaction in male mice. Eur J Neurosci. 2011; 34(3): 469–77.

43. Flanagan LM, Pfaus JG, Pfaff DW, McEwen BS. Induction of FOS immunoreactivity in oxytocin neurons after sexual activity in female rats. Neuroendocrinology. 1993; 58(3): 352–8.

44. Cho MM, DeVries AC, Williams JR, Carter CS. The effects of oxytocin and vasopressin on partner preferences in male and female prairie voles (Microtus ochrogaster). Behav Neurosci. 1999; 113(5): 1071–9.

45. Song Z, Larkin TE, Malley MO, Albers HE. Oxytocin (OT) and arginine-vasopressin (AVP) act on OT receptors and not AVP V1a receptors to enhance social recognition in adult Syrian hamsters (Mesocricetus auratus). Horm Behav. 2016; 8120–7.

46. Gabor CS, Phan A, Clipperton-Allen AE, Kavaliers M, Choleris E. Interplay of oxytocin, vasopressin, and sex hormones in the regulation of social recognition. Behav Neurosci. 2012; 126(1): 97–109.

47. Madlon-Kay S, Montague MJ, Brent LJN, Ellis S, Zhong B, Snyder-Mackler N, Horvath JE, Skene JHP, Platt ML. Weak effects of common genetic variation in oxytocin and vasopressin receptor genes on rhesus macaque social behavior. Am J Primatol. 2018; 80(10): e22873.

48. Wacker D, Ludwig M. The role of vasopressin in olfactory and visual processing. Cell Tissue Res. 2019; 375(1): 201–15.

49. Vaz RP, Cardoso A, Sá SI, Pereira PA, Madeira MD. The integrity of the nucleus of the lateral olfactory tract is essential for the normal functioning of the olfactory system. Brain Struct Funct. 2017; 222(8): 3615–37.

50. Krause EG, de Kloet AD, Flak JN, Smeltzer MD, Solomon MB, Evanson NK, Woods SC, Sakai RR, Herman JP. Hydration state controls stress responsiveness and social behavior. J Neurosci. 2011; 31(14): 5470–6.

51. de Bono M, Bargmann CI. Natural variation in a neuropeptide Y receptor homolog modifies social behavior and food response in C. elegans. Cell. 1998; 94(5): 679–89.

52. Hernandez-Perez OR, Hernandez VS, Nava-Kopp AT, Barrio RA, Seifi M, Swinny JD, Eiden LE, Zhang L. A Synaptically Connected Hypothalamic Magnocellular Vasopressin-Locus Coeruleus Neuronal Circuit and Its Plasticity in Response to Emotional and Physiological Stress. Front Neurosci. 2019; 13196.

